# A role for gut microbiota in m^6^A epitranscriptomic mRNA modifications in different host tissues

**DOI:** 10.1101/504266

**Authors:** Sabrina Jabs, Christophe Becavin, Marie-Anne Nahori, Vincent Guérineau, David Touboul, Amine Ghozlane, Marie-Agnès Dillies, Pascale Cossart

## Abstract

The intestinal microbiota modulates host physiology and gene expression via mechanisms that are not fully understood. A recently discovered layer of gene expression regulation is N6-methyladenosine (m^6^A) modification of mRNA. To unveil if this epitranscriptomic mark in part mediates the impact of the gut microbiota on the host, we analyzed m^6^A-modifications in transcripts of mice displaying either a conventional, or a modified, or no gut flora. We discovered that the microbiota has a strong influence on m^6^A-modifications in the cecum, and also, albeit to a lesser extent, in the liver. We furthermore show that a single commensal bacterium, *Akkermansia muciniphila*, can affect specific m^6^A modifications. Together, we report here epitranscriptomic modifications as an unexpected level of interaction in the complex interplay between commensal bacteria and their host.

## Introduction

Posttranscriptional mRNA modifications, most notably m^6^A (Dominissini et al., 2012; Meyer et al., 2012), have recently been shown to contribute to the regulation of mRNA fate by affecting mRNA stability, splicing events or the initiation of translation (Peer et al., 2017). mRNA can be methylated by RNA-methyltransferases, in specific positions that are mainly located at the 3’ untranslated regions (UTR) and the coding sequence (CDS) of the transcript, utilizing S-adenosyl methionine (SAM) as a methyl donor. METTL3 in complex with METTL14 is the most important m^6^A-modifying enzyme (Liu et al., 2013), but for specific transcripts, METTL16 has been proposed to act as an additional N6-adenosine-methyltransferase (Pendleton et al., 2017; Warda et al., 2017). m^6^A modifications can be removed by the demethylases AlkbH5 and FTO. The latter has been shown to be associated with obesity in humans and mice (Church et al., 2009; Fischer et al., 2009; Frayling et al., 2007), thus representing important regulator of host metabolism. m^6^A modification of mRNA has been shown to be important in embryonic stem cell and immune cell differentiation (Frye and Blanco, 2016; Geula et al., 2015; Li et al., 2017), neurogenesis (Yoon et al., 2017), stress responses (Meyer et al., 2015), the circadian rhythm (Fustin et al., 2013), and viral infection (Gokhale et al., 2016; Lichinchi et al., 2016a; Lichinchi et al., 2016b; Tan and Gao, 2018; Tirumuru et al., 2016). Whether bacteria influence m^6^A modifications is unknown. Commensal bacteria, in particular the gut microbiota, have profound effects on host physiology, such as host metabolism, intestinal morphology, the development of the immune system, and even behavior (Sommer and Bäckhed, 2016). Gut microbial metabolites and fermentation products, e.g. short chain fatty acids (SCFA), sphingolipids and tryptophan metabolites have been shown to partially mediate the influence of gut commensals on their host (Agus et al., 2018; Heaver et al., 2018; Koh et al., 2016). However, many aspects of gut-microbiota-host interactions still remain obscure. By using methylated RNA-immunoprecipitation and sequencing (MeRIP-Seq), we set out to determine if the influence of the microbiota on host is mediated by alteration of m^6^A mRNA modification profiles.

## Results and discussion

### m^6^A modification profiles in cecum and liver

For our study we focused on two different organs: the cecum, which is in close contact with the gut microbiota, and the liver, whose gene expression is also known to be influenced by commensal bacteria (Manes et al., 2017), and used mice with different gut florae (see below). Overall, we detected 86,418 methylation sites in anti-m^6^A-immunoprecipitates from murine cecum and 54,319 methylation sites in anti-m^6^A-immunoprecipitates from liver. 39,921 methylation sites were common between the two organs (Figure S1A), and 81% and 84% of the sites we detected in the cecum and liver, respectively, are described in the Methyl Transcriptome Data Base (Liu et al., 2018) (Figure S1A), indicating that we identified *bona fide* methylation sites. The m^6^A marks that we identified were mostly present in the CDS and 3’UTR of mRNA, but also in the 5’ UTR (Fig. S1B), in agreement with previous studies (Dominissini et al., 2012; Meyer et al., 2012).

### Gut flora controls m^6^A modifications in mouse cecum

We first compared m^6^A marks in the cecal transcripts of conventional (CONV) and germ-free (GF) mice (Figure 1A) and found 771 m^6^A sites on 431 transcripts to be differentially methylated. Principal component analysis (PCA) of m^6^A sites revealed a clear separation between methylation sites of CONV and GF mice in the cecum (Figure 1B). Furthermore, the heat map with hierarchical clustering of methylation sites (Figure 1C), which considers the 500 differentially methylated sites with the highest variability, showed a strong separation of CONV and GF cecal transcripts. Importantly, the transcript expression levels of the majority (63%) of differentially methylated m^6^A sites were not significantly changed (Figure 1D), excluding that the differences in methylation between the biological conditions are due to altered transcript levels, as reported in other cases (Schwartz et al., 2014). To test whether differential methylation was mediated by the gut microbiota, we colonized GF mice with the flora of CONV mice (ex-GF). After four weeks, these mice exhibited the same patterns of the most abundant gut bacterial genera (e.g. *Lachnospiraceae, Allistipes, Bacteroides, Prevotellaceae, Akkermansia*) as CONV mice (Figure 1A, Figure S2A). m^6^A marks in the ceca of ex-GF mice clustered with the methylation sites of CONV mice in PCA and the heatmap, and only four m^6^A sites on four transcripts were differentially methylated when comparing CONV and ex-GF mice (Figure 1D), demonstrating unambiguously that the gut microbiota mediates differential methylation.

**Fig. 1:**
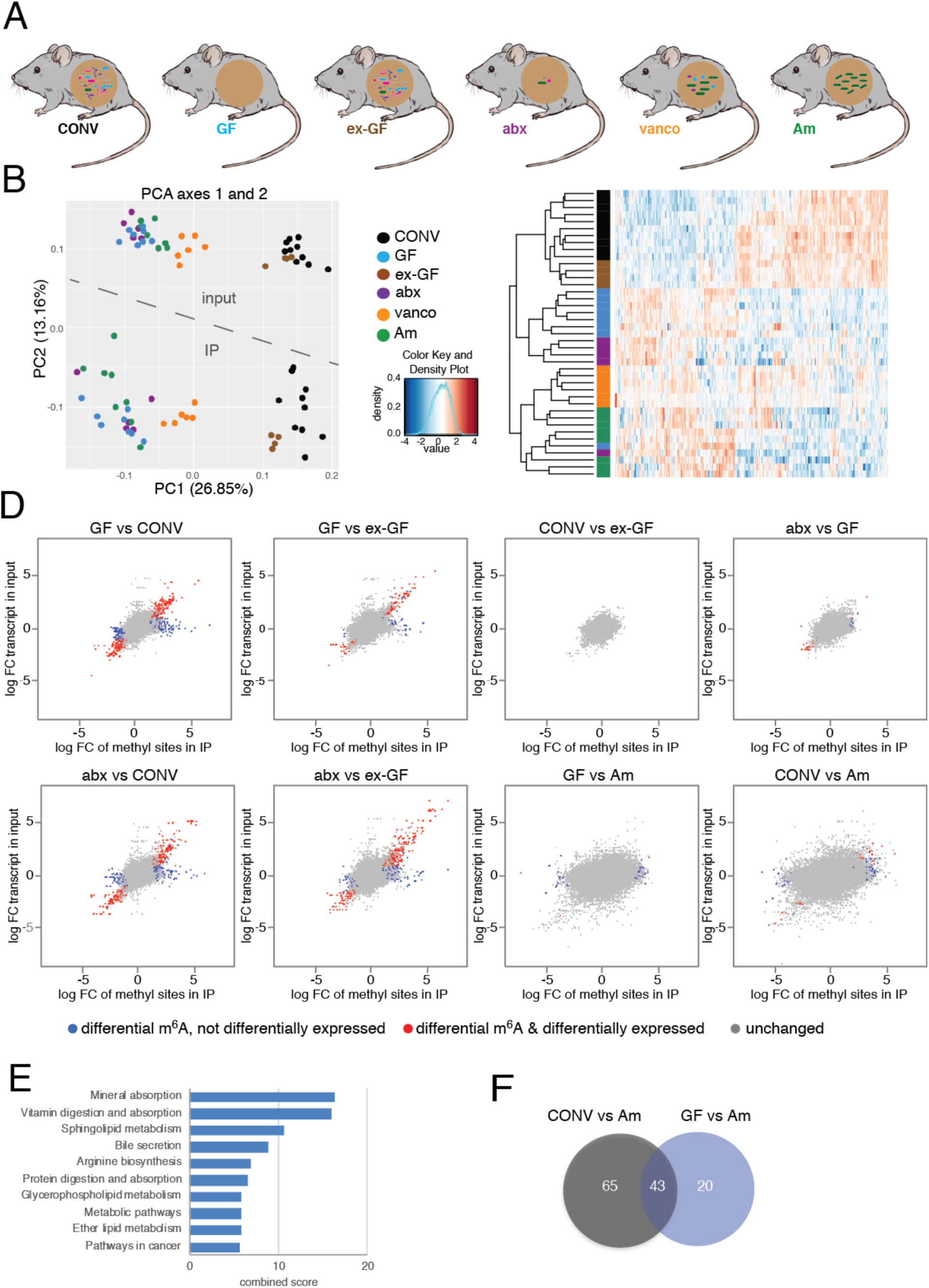
Microbiota influences m^6^A modification in mouse cecum. **A**, The six mouse models analyzed: CONV, conventionally raised mouse (n=10); GF, germ-free mouse (n=8); ex-GF, GF mouse colonized with the gut flora of CONV mice (n=4); abx, CONV mice whose gut flora has been depleted by antibiotics treatment (n=5); vanco, vancomycin/amphotericinB-treated mice (n=7); Am, *A*.*muciniphila*-mono-colonized mice (n=8). **B**, principal component analysis of normalized read counts of the 500 most variable differentially methylated sites and corresponding regions in input; **C**, heatmap of normalized read counts of the 500 most variable differentially methylated sites; **D,** m^6^A-sites found to be differentially methylated compared to differential expression of transcripts in indicated mice. Differentially methylated sites that are also differentially expressed on transcript level are in red, differentially methylated sites that are unchanged on transcript level, are in blue. m^6^A sites that were not significantly changed are shown in grey. Cut-offs for differential expression are log fold change (FC) −1 to 1 and Benjamini-Hochberg-corrected p-values <0.05. **E,** KEGG pathway analysis of differentially methylated transcripts between CONV and GF mice performed using the Enrichr tool (Chen et al., 2013; Kuleshov et al., 2016). **F**, Venn diagram of transcripts differentially methylated between Am and CONV, and Am and GF, respectively.

To confirm these results, we analyzed mice treated with a mix of several antibiotics (vancomycin, metronidazole, neomycin, ampicillin and the antifungal amphotericin B) for three weeks (abx mice), which resulted in an efficient depletion of the gut flora with only few genera still detectable by 16S sequencing (e.g. *Akkermansia, Escherichia/Shigella, Parasutterella, Lactobacillus* and *Lachnospiraceae*) after the treatment (Figure 1A, Figure S2B, C). m^6^A-modified marks in transcripts from abx cecum were found in clusters mixed with (Figure 1B) and closely related to (Figure 1C) those in GF mice, demonstrating that the impact of a conventional flora on m^6^A can be suppressed by antibiotics treatment. Similar to the situation in GF mice compared to CONV mice, differentially methylated transcripts in abx mice, were mostly unchanged on transcript level (Figure 1D). A number of m^6^A sites (76) were still differentially methylated between abx and GF mice (Figure 1D), which can be explained by the influence of the few commensals still present after abx treatment (Figure S2C). Taken together, these results establish that the gut microbiota regulates posttranscriptional mRNA modifications in the host in addition to well-known effects on transcription and protein expression.

To gain further insight into the bacterial species that may be involved in the regulation of differential m^6^A modifications, we treated mice with only vancomycin, to which we equally added the antifungal amphotericin B (vanco mice). This treatment resulted in a strong enrichment of many of the genera that persisted in abx mice (*Akkermansia, Escherichia/Shigella, Lactobacillus, Lachnospiraceae, Parasutterella*; Figure S2B). In the PCA, m^6^A sites in vanco mice formed a cluster distinct from the GF/abx and CONV/ex-GF mice clusters (Figure 1B). As expected, vanco mice exhibited less differentially methylated sites compared to CONV mice (60 sites) than GF (771) or abx (657) mice, respectively, compared to CONV mice (Figure 1D, Figure S3). These data suggest that the persisting commensals in vanco mice are involved in the gut microbiota-mediated regulation of m^6^A modification patterns observed in CONV cecum.

### *Akkermansia muciniphila* induces differential m^6^A modifications in the cecum

We proceeded to testing if differential methylation can also be induced by mono-association of GF mice with a single bacterial species. We chose *A*.*muciniphila*, which was still present or the most enriched, respectively, in abx and vanco mice, and has been shown to be an important commensal influencing host physiology (Derrien et al., 2017). We successfully mono-associated GF mice with *A*.*muciniphila* (Am; Figure 1A; Figure S1D) and found that differentially methylated transcripts in ceca of Am mice were found in clusters with those of GF and abx mice in the PCA (Fig. 1B). However, the heatmap with hierarchical clustering revealed a pattern of m^6^A sites most closely related to the methylation sites in vanco mice (Figure 1C). In total, we found 119 m^6^A sites on 108 transcripts to be differentially methylated between CONV and Am mice. 63 sites on 63 transcripts were differentially methylated between GF and Am mice, suggesting that *A*.*muciniphila* is indeed able to induce differential expression of m^6^A-modified marks in GF mice. Interestingly, there was a substantial overlap of common differential m^6^A-modifications between Am and CONV and Am and GF (Figure 1F). These results let us conclude that there are differentially methylated transcripts specifically affected by *A*.*muciniphila*.

### The gut flora regulates differential methylation of transcripts involved in metabolic pathways and amino acid synthesis

To decipher the cellular functions affected by microbiota-dependent modifications, we performed pathway analyzes on the list of differentially methylated transcripts between CONV and GF cecum. KEGG pathway analysis revealed an enrichment of differentially methylated transcripts involved in degradation and absorption processes (minerals, vitamins, proteins), metabolic pathways of lipids, and amino acid synthesis (Figure 1E). Several of these processes, e.g. synthesis of amino acids, and lipid metabolism, have previously been shown to be influenced by the microbiota (Bäckhed et al., 2004; Mardinoglu et al., 2015). However, our results establish posttranscriptional modifications as an additional and so far unexpected layer of gene expression regulation in these pathways.

### Gut microbiota also affects m^6^A modifications in the liver

To examine whether the influence of the gut microbiota on m^6^A mRNA modifications is specific to the cecum, and as transcript expression in the liver is also known to be affected by the gut microbiota, we performed differential MeRIP-Seq analysis of transcripts from liver of CONV and GF mice (Figure 2A). In total, we detected 230 sites on 106 transcripts to be differentially methylated in liver, revealing an influence of the microbiota on m^6^A modifications in this organ, albeit less important than in the cecum. PCA (Figure 2B) and heat maps (Figure 2C) showed clusters of m^6^A sites in the liver of CONV mice that were clearly different from m^6^A sites on transcripts from GF liver. Similar to what we observed in the cecum, the majority (63%) of the few differentially methylated transcripts are altered without changes on transcript level (Figure 2D). Together, these results show that the regulation of host m^6^A by the gut microbiota is not restricted to the intestine and represents a general mechanism that influences host physiology.

**Fig. 2:**
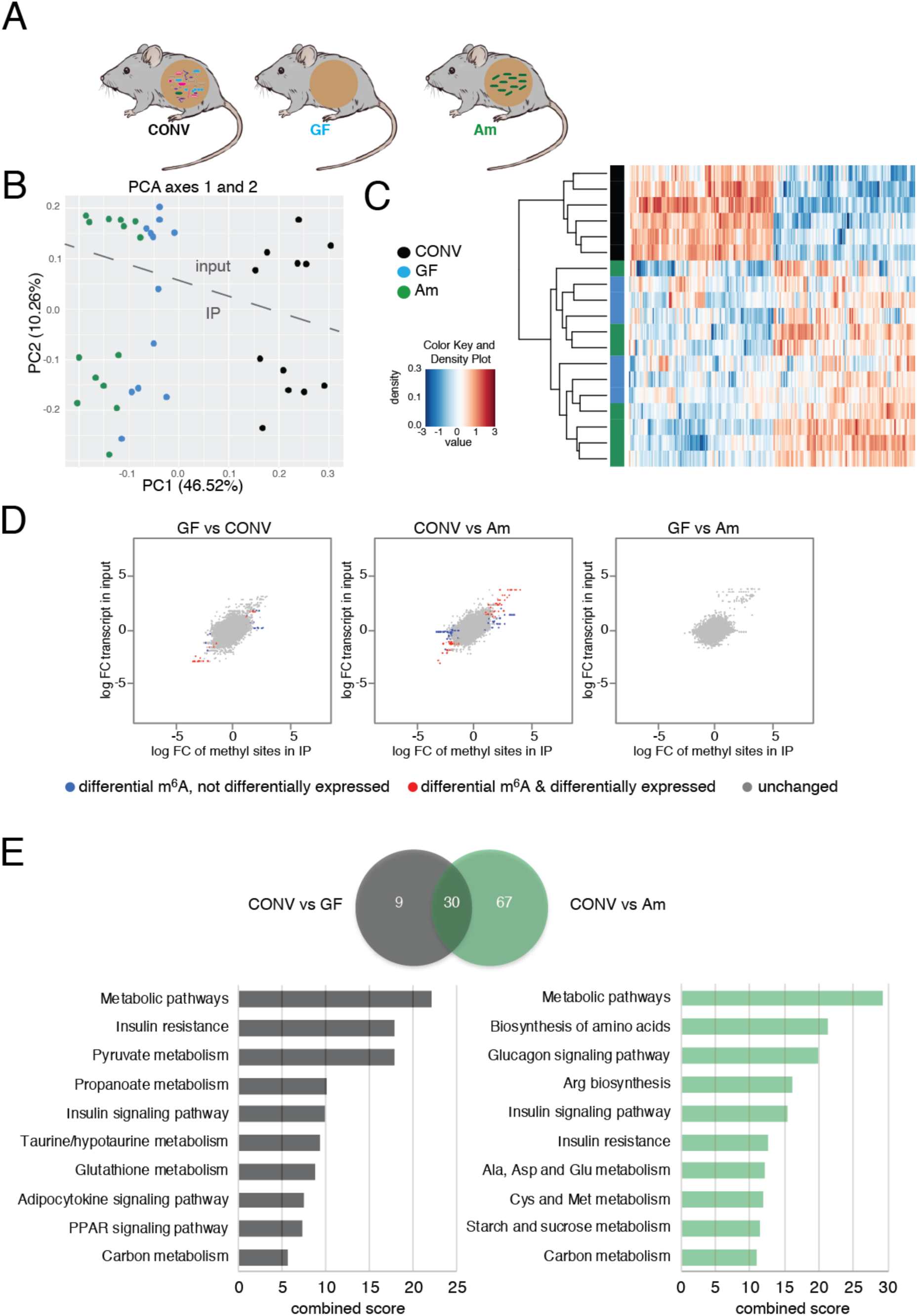
Microbiota influences m^6^A modification in mouse liver. **A**, The three mouse models analyzed: CONV, conventionally raised mouse (n=6); GF, germ-free mouse (n=6); Am, *A*.*muciniphila*-mono-colonized mice (n=7) Tissues were derived from 2 independent experiments. **B**, principal component analysis of normalized read counts of all differentially methylated sites and corresponding regions in input; **C**, heatmap of normalized read counts of all differentially methylated sites; **D**, m^6^A-sites found to be differentially methylated compared to differential expression of transcripts in indicated mouse models. Differentially methylated sites that are also differentially expressed on transcript level are in red, differentially methylated sites that are unchanged on transcript level, are in blue. Methylation sites that were not significantly changed are shown in grey. Cut-offs for differential expression are log fold change (FC) −1 to 1 and Benjamini-Hochberg-corrected p-values <0.05. **E**, Venn diagram comparing differentially methylated transcripts between CONV vs GF and CONV vs Am. KEGG pathway analysis of differentially methylated transcripts between CONV and GF mice and transcripts differentially methylated between CONV and Am using the Enrichr tool (Chen et al., 2013; Kuleshov et al., 2016).

### *A. muciniphila* controls differential methylation of transcripts involved in amino acid metabolism in the liver

Colonization of GF mice with *A*.*muciniphila* revealed m^6^A-modified transcripts in liver that were also differentially modified in GF mice in PCA and heatmap (Figure 2B, C). However, the number of significantly differentially methylated transcripts between CONV and Am mice in the liver is higher than between CONV and GF mice, suggesting an important impact of *A*.*muciniphila* on m^6^A-modifications in this organ (Fig. 2D). KEGG analysis demonstrated that in GF and Am mice, respectively, compared to CONV mice, metabolic pathways were the most affected by differential methylation, including transcripts involved in glucose and insulin signaling, which have previously been shown to be influenced by the gut microbiota (Nicholson et al., 2012; Schroeder and Bäckhed, 2016; Shapiro et al., 2018). However, transcripts involved in amino acid synthesis and metabolism (Arg, Ala, Asp, Glu, Cys, and Met) were more enriched among differentially methylated transcripts between CONV and Am, indicating an involvement of *A*.*muciniphila* in the regulation of these pathways (Figure 2E, Table S4).

### Differentially methylated transcripts are more frequently modified in the 5’UTR of mRNAs

In order to identify possible common features of the differentially methylated transcripts, we analyzed the positions of all differentially methylated sites in both tissues using the GUITAR package (Liu et al., 2018) and found that the position of methylation sites in transcripts with unchanged expression levels, although still present in the CDS and 3’UTR, was shifted towards the 5’ UTR compared to the total detected methylation sites in the cecum (Figure 3A). This effect was even more pronounced in the liver, where all differentially methylated sites were more frequently detected in the 5’UTR (Figure 3B). It is still unclear how specific subsets of transcripts can be methylated in a microbiota-dependent manner. However, the increased proportion of m^6^A-marks present in the 5’UTR of differentially methylated transcripts may be explained by the presence of a specific motif or structural arrangement favoring recognition by a specific methyltransferase complex, or accessory proteins, thus conferring specificity to distinct subsets of transcripts.

**Fig. 3:**
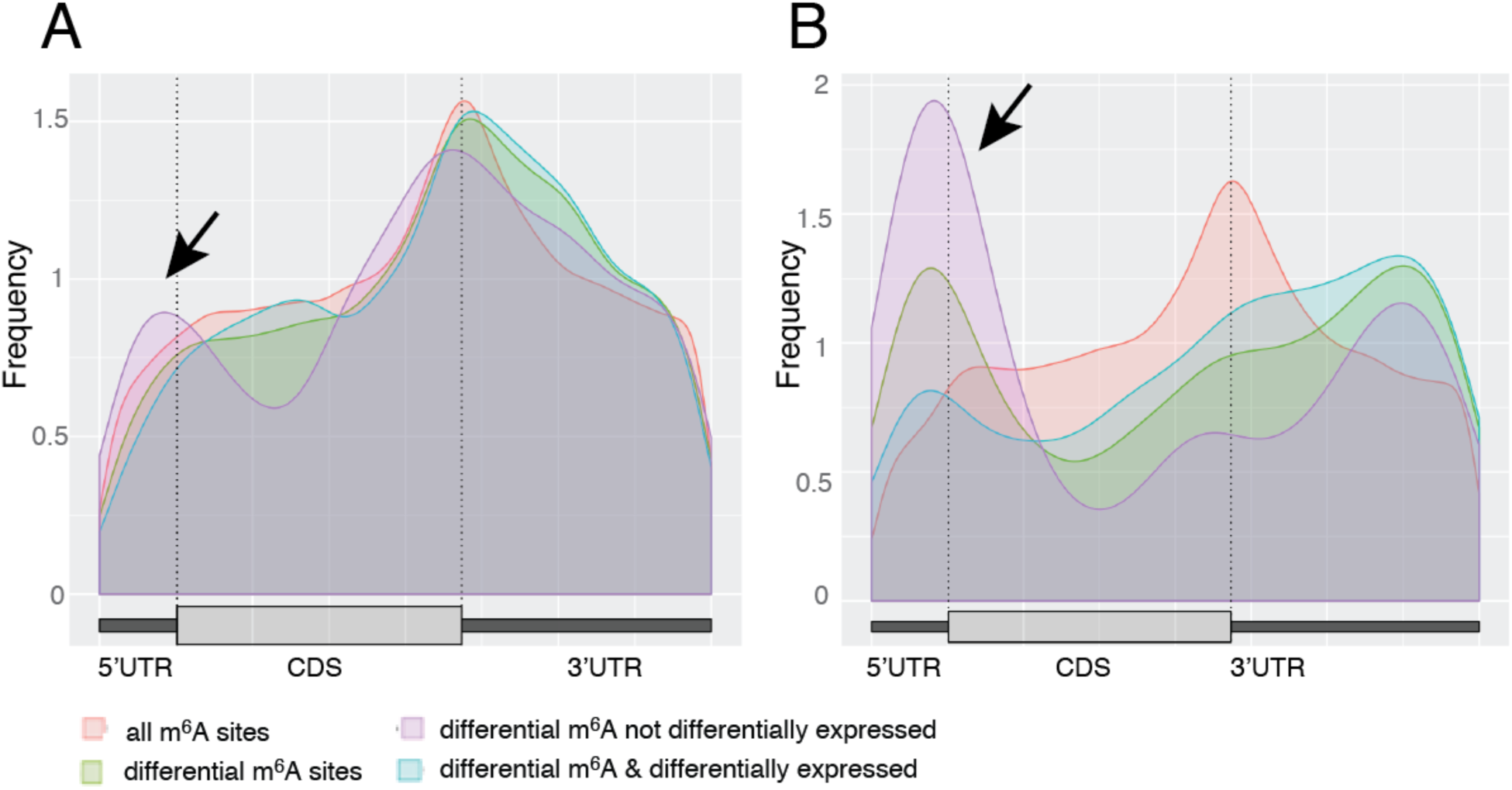
Analysis of specificity of microbiota-dependent mRNA methylation. GUITAR (Liu et al., 2018) plots for distribution of all detected m^6^A sites from **A**, cecum and **B**, liver. All detected m^6^A sites are depicted in red, differentially methylated sites are in green. Differentially methylated sites with altered transcript expression are in blue, and differentially methylated sites with unchanged transcript expression are in purple. UTR, untranslated region; CDS, coding sequence. Arrow indicates the shift of differentially methylated sites that are not differentially expressed towards the 5’UTR.

### Differential methylation of Mat2a links gut commensal metabolism and host mRNA methylation

The precise mechanisms of regulation for m^6^A-methyltransferases are not known, but it is clear that their activity requires the availability of the main methyl donor, S-adenosylmethionine (SAM). We found the transcript of the SAM-producing enzyme Mat2a to be less methylated in the cecum of GF, abx and Am mice compared to CONV, ex-GF, and vanco mice (Fig. 4A, B, Fig. S4, Table S1). Interestingly, in cultured cells Mat2a expression has recently been shown to be regulated by m^6^A modification (Pendleton et al., 2017; Shima et al., 2017): high concentrations of SAM cause increased m^6^A modification of Mat2a mRNA in its 3’UTR, which leads to an enhanced degradation of the transcript, thereby down-regulating Mat2a protein expression (Pendleton et al., 2017; Shima et al., 2017), and decreasing intracellular SAM synthesis. Since SAM levels are influenced by nutrient uptake and the gut microbiota (Krautkramer et al., 2016), the lower methylation of Mat2a mRNA in GF, abx and Am mice may reflect the lower supply of SAM or metabolites required for its synthesis from the intestinal content of these mice. Mat2a is not expressed in the liver, and the liver-specific isoform Mat1a has not been shown to be regulated by SAM-dependent m^6^A-modification of the corresponding transcript. In agreement with this observation, we found that the majority of the 32 m^6^A sites we detected in Mat1a was not influenced by the microbiota (Table S2).

**Fig. 4:**
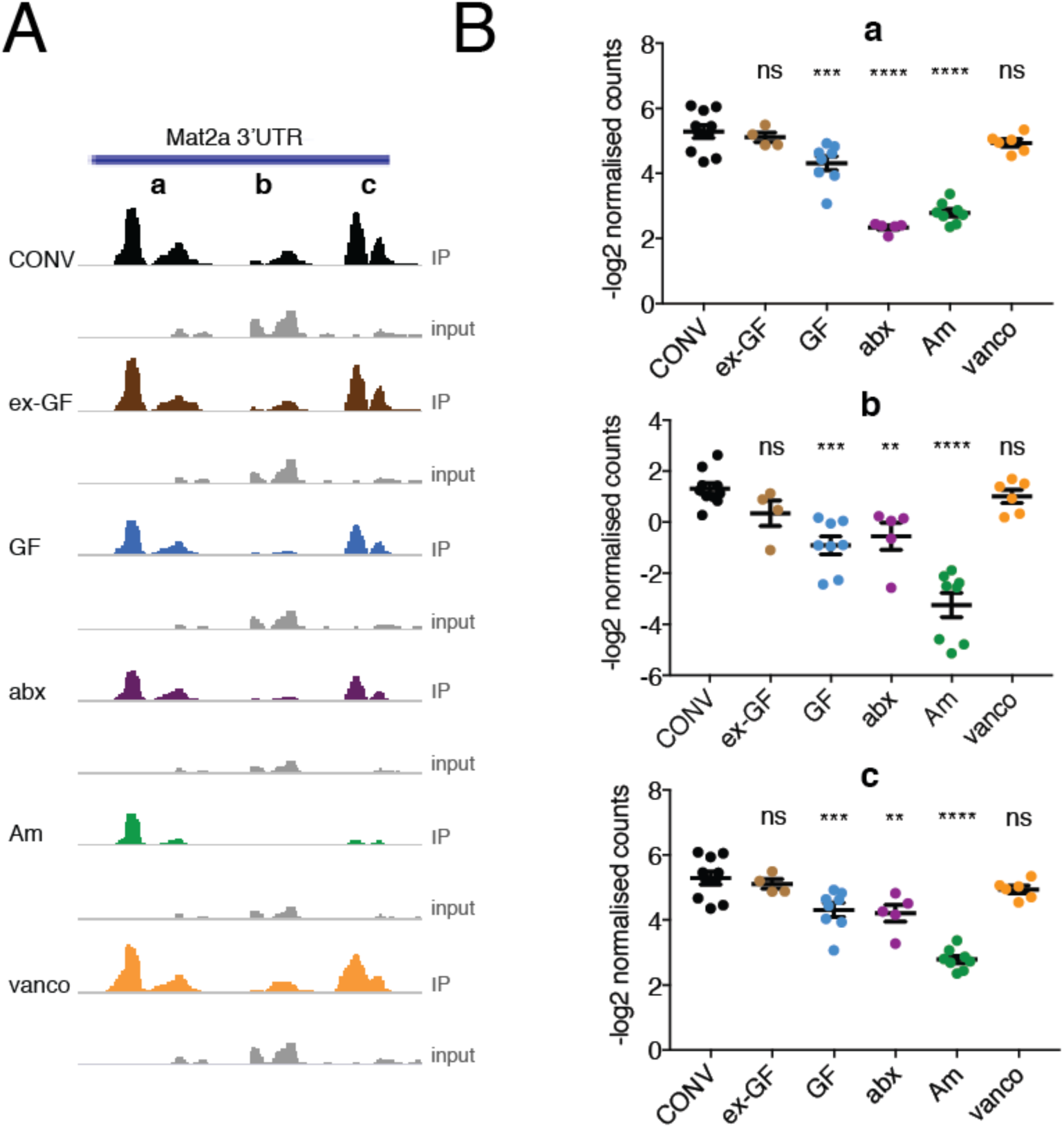
Differential methylation of Mat2a mRNA in the 3’UTR. **A,** Mean of read per million normalized coverage (RPM) in detected methylation sites from anti-m^6^A immunoprecipitates and input in the 3’UTR of the Mat2a transcript in cecum (CONV, n=10; GF, n=8; ex-GF, n=4; abx, n=5; Am, n=8; vanco, n=7) visualized using IGV; **B**, quantification of select Mat2a sites (**a**,**b**,**c** from **A**) as −log2 normalized read counts. Ordinary one-way ANOVA for multiple comparisons was performed. * p-value < 0.05; ** p-value< 0.01; *** p-value < 0.005; **** p-value < 0.001; ns, non-significant.

Taken together, we identified epitranscriptomic modifications as a new mechanism of commensal-host-interaction, setting the ground for future studies on the regulation of a microbiota-directed methylation machinery and the translational consequences of this regulatory process.

## Acknowledgments

The authors thank Marion Bérard and the Gnotobiology platform of the Pasteur Institute for support in conducting gnotobiology experiments; the Pasteur Biomics platforms for Transcriptomics and Epigenetics and Metagenomics, headed by Jean-Yves Coppée and Sean Kennedy, respectively, for providing RNA sequencing and 16S DNA sequencing services; Carla Saleh and Olivier Dussurget for helpful discussions; Hugo Varet (Institut Pasteur Bionformatics and Statistics Hub) for advice concerning statistical analysis.

## Funding

This work was supported by grants to P.C.: European Research Council (ERC) Advanced Grant BacCellEpi (670823), ANR Investissement d’Avenir Programme (10-LABX-62-IBEID), ERANET-Infect-ERA PROANTILIS (ANR-13-IFEC-0004-02) and the Fondation Le Roch-Les Mousquetaires. S.J. was supported by a Danone fellowship. P.C. is a Senior International Research Scholar of the Howard Hughes Medical Institute.

## Author contributions

S.J. designed and conducted most of the experiments, analyzed data and wrote the manuscript; C.B. performed bioinformatics analysis; M.A.N. helped with mouse experiments; V.G. and D.T. performed LC-HRMS experiments; A.G. analyzed 16S DNA sequencing data; M.A.D. supervised bioinformatics and performed statistics analyzes; P.C. designed the overall study and wrote the manuscript.

## Competing interests

The authors declare no competing interests.

## CONTACT FOR REAGENT AND RESOURCE SHARING

Further information and requests for resources and reagents should be directed to and will be fulfilled by the Lead Contacts Pascale Cossart and Sabrina Jabs.

## MATERIAL and METHODS

### Bacterial strains and culture conditions

*Akkermansia muciniphila* (ATCC BA-835), obtained from the Biological Resource Center of Institut Pasteur (CRBIP), was cultured in Brain Heart Infusion (BHI) supplemented with 8mM L-Cysteine hydrochloride, 0.2% NaHCO3, and 0.025% Hemin in an anaerobic atmosphere using Oxoid™ AnaeroGen™ 2.5L gas packs (Thermo Fisher) at 37°C.

### Animal experiments

All animal experiments were approved by the committee on animal experimentation of the Institut Pasteur and by the French Ministry of Agriculture.

### Mice

C57BL/6J mice were purchased from Charles River. Germ-free mice generated from C57BL/6J mice were obtained either from CNRS TAAM UPS44 Orléans, France, or from the Gnotobiology Platform of the Institut Pasteur and kept in isolators. Conventional mice were kept in specific pathogen-free conditions and all the mice used were female. Mice were housed in 10h (dark)/ 14h (light) cycles.

### Colonization of germ-free mice

For generating ex-GF mice, 10-12 fecal pellets were collected from 4 CONV mice housed in the same cage for 2 weeks, resuspended, and added to the drinking water of GF mice. This procedure was repeated on 3 consecutive days. On Day 4, cecal content of the 4 CONV mice was collected, resuspended in 10 mL of PBS, of which 0.2 mL was administered to the pre-colonized mice by oral gavage. Colonization efficiency was determined in the cecal content of mice after 4 weeks. For mono-colonization experiments, germ-free mice were inoculated once with 10^9^ colony-forming units (CFU) of *A*. *muciniphila* and maintained for four weeks. Mono-colonization was monitored by *A*. *muciniphila*-specific PCR (Collado et al., 2007) and controlled by 16S DNA sequencing in cecal content (not shown).

### Antibiotic treatment of mice

For depletion of the gut flora, conventional C57BL/6J mice were treated with antibiotics as described previously (Reikvam et al., 2011). In brief, after oral treatment with amphotericin B (0.1 mg/mL; Sigma Aldrich) for 2-3 days, mice were treated with a solution consisting of 10 mg/mL ampicillin, 5 mg/mL vancomycin 10 mg/mL neomycin, 10 mg/mL metronidazol and 0.1 mg/ml amphotericin-B (all Sigma Aldrich) *per os* every 12h using a gavage volume of approximately 10 mL/kg body weight for 21 days. The depletion was controlled for by quantitative PCR as described (Reikvam et al., 2011) and the identity of residual bacterial genera determined by 16S DNA sequencing. For enrichment of a small number of genera, mice were treated as above with only 5 mg/mL vancomycin and 0.1 mg/ml amphotericin-B. Tissues used for the analysis were derived from 2-3 independent experiments.

### RNA preparation and ribodepletion

Total RNAs were prepared using the RNeasy maxi kit (Qiagen) and quality-controlled using RNA 6000 Nano assay (Agilent). Ribodepletion was performed using RiboMinus Transcriptome Isolation Kit (Human/Mouse; Thermo Fisher) and controlled using RNA 6000 Pico assay (Agilent). We performed preliminary experiments to determine absolute concentrations of select mRNA modifications by liquid chromatography coupled to high resolution mass spectrometry (LC-HRMS, see below). We could not detect significant changes in nucleoside concentrations using this method (Figure S5).

### LC-HRMS

Ribodepleted RNAs were desalted using Microcon YM10 columns (Millipore) and subjected to Nuclease P1 digestion in 50mM ammonium acetate (pH 7) in the presence of Antarctic phosphatase (New England Biolabs) as described previously (Suzuki et al., 2007). Nucleoside composition was analyzed by narrow bore HPLC using a U-3000 HPLC system (Thermo-Fisher). An Accucore RP-MS (2.1 mm X 100 mm, 2.6 µm particle) column (Thermo-Fisher) was used at a flow rate of 200 µl/min at a temperature of 30°C. Mobile phases used were 5 mM ammonium acetate, pH 5.3 (Buffer A) and 40% aqueous acetonitrile (Buffer B). A multilinear gradient was used with only minor modifications from that described previously (Pomerantz and McCloskey, 1990).

An LTQ Orbitrap™ mass spectrometer (Thermo Fisher Scientific) equipped with an electrospray ion source was used for the LC/MS identification and quantification of nucleosides. Mass spectra were recorded in the positive ion mode over an *m/z* range of 100-1000 with a capillary temperature of 300°C, spray voltage of 4.5 kV and sheath gas, auxiliary gas and sweep gas of 40, 12 and 7 arbitrary units, respectively. Calibration curves were generated using a mixture of synthetic standards of Adenosine (A) and Cytidine (C) (Sigma-Aldrich), m^6^A, and m^1^A, m^5^C (TCI Europe), and N^6^, 2’-O-dimethyladenosine (m^6^Am; Berry&Associates) in the ranges of 20-625 injected fmol for m^1^A, m^6^A, m^6^Am and m^5^C and 5-250 injected pmol for A and C. Each calibration point was injected in triplicate. Extracted Ion Chromatograms (EIC) of base peaks of the following masses: A (*m/z* 268.08-268.12), C (*m/z* 244.08-244.11), m^1^A and m^6^A (*m/z* 282.10-282.13), m^6^Am (*m/z* 296.12-296.15), m^5^C (*m/z* 258.09-258.12), were used for quantification. In all cases, coefficients of variations for peak areas were always below 15%. Experimental data (peak area *versus* injected quantity) were fitted with a linear regression model for each compound leading to coefficient of determination (R^2^) values better than 0.997. Accuracies were calculated for each calibration point and were always better than 15%.

### Immunoprecipitation of m^6^A-methylated mRNA

Immunoprecipitation was performed as previously described (Schwartz et al., 2013). Briefly, 3 μg of rabbit anti m^6^A antibody (Synaptic Systems) were bound to 25 μl washed Protein G Dynabeads (Thermo Fisher) in immunoprecipitation buffer (1x IPP; 150 mM NaCl, 0.1% NP-40, 10 mM Tris-Cl, pH 7.4) for 30 min at room temperature (RT) and washed twice in 1x IPP. Total (liver) or ribodepleted (cecum) RNA was fragmented using the NEBNext Magnesium RNA fragmentation module (New England Biolabs), purified by ethanol precipitation and quality-controlled using the RNA 6000 pico assay (Agilent). Equal amounts of RNA (5 μg for ribodepleted and 200 μg for total RNA) were denatured for 2 min at 70°C and adjusted to 1x IPP concentration using 2x IPP. RNA was added to the antibody-bound beads and incubated for 2h at 4°C in the presence of murine RNase inhibitor (New England Biolabs). The bound RNA was washed twice with 1x IPP, twice with low salt IPP buffer (50 mM NaCl, 0.1% NP40, 10 mM Tris-Cl pH 7.4), twice with high salt IPP buffer (500 mM NaCl, 0.1% NP40, 10 mM Tris-Cl pH 7.4) and once more in 1x IPP. RNA was eluted using 30 μl of buffer RLT (Qiagen). 20 μl of MyOne Silane Dynabeads (Thermo Fisher) were washed with RLT and resuspended in 30 μl of RLT. Eluted RNA was bound to the beads in the presence of 35 μl of absolute ethanol, washed twice in 70% ethanol and eluted in 100 μl of H_2_0. RNA was purified and concentrated using RNA clean & concentrator (Zymo research) before proceeding to library preparation.

### Library Preparation and sequencing

150 ng of input RNA and the immunoprecipitated RNAs were dephosphorylated using Antarctic phosphatase and subjected to T4 PNK treatment (both New England Biolabs). Directional m^6^A RNA-seq libraries were prepared using NEBNext Multiplex Small RNA Library Prep Set for Illumina (New England Biolabs) according to the manufacturer’s instructions. Libraries were sequenced on an Illumina HiSeq 2500 platform generating single end reads (65bp).

### MeRIP-Seq processing

The mouse mm10 genome and list of transcripts were downloaded from Gencode (*Mus Musculus* VM13(Mudge and Harrow, 2015)). Only the 21968 ‘protein_coding’ genes were kept for the analysis. After the sequencing of every MeRIP-Seq (IP dataset) and RNASeq (Input dataset) the resulting reads were trimmed (AlienTrimmer 0.4.0(Criscuolo and Brisse, 2013), default parameters). They were mapped on mouse mm10 genome using STAR mapper 2.5.0a *(--sjdbOverhang 100* parameter) (Dobin et al., 2013). Mapping files were filtered to keep uniquely mapped reads using SAMtools 0.1.19 (*samtools view -b –q 1* parameters) (Li et al., 2009), and saved to BAM files after indexation. The quality of the sequencing and mapping was assessed using FastQC 0.10.1 and MultiQC 0.7 (Ewels et al., 2016). Gene expression was calculated with HTSeq 0.9.1(-s no -m union --nonunique all parameters) (Anders et al., 2015).

### m^6^A site detection

The original three reference papers on MeRIP-Seq analysis have used three different workflows for m^6^a modification site detection (Fisher test (Meyer et al., 2012), MACS2 software (Dominissini et al., 2013), and Peak Over Input technique (Schwartz et al., 2013)). Each of these techniques has a certain bias and identifies different types of methylation sites. We implemented all of and developed our own technique using the fold of Reads Per Million (RPMF). We prepared the peak detection by first generating windows of 100bp overlapping in their middle using all 21,968 ‘protein_coding’ genes, windows of 100bp overlapping in their middle were generated. The total number of reads per window was calculated for each dataset. The number of reads for each window in IP and Input was determined using HTSeq 0.9.1 (Anders et al., 2015)*(-s no -m union --nonunique all* parameters). Only the windows with coverage higher than 10 reads in IP datasets were kept. Fisher, POI, and RPMF techniques were then run on these windows to assess for the presence of methylation sites.

The Fisher exact rank test was applied on each of the 100bp windows. The p-values were corrected with Benjamini-Hochberg multiple testing correction (Benjamini, 1995). Only peaks with a corrected p-value < 0.001 were kept for Fisher analysis. The reads per million (RPM) for each window in IP and Input was calculated. The reads per million-fold (RPMF) was determined by subtraction of RPM of each window in the IP dataset and RPM in the Input. Only the peaks with RPMF > 10 were then kept. We prepared the POI peak detection by first calculating the raw coverage on each dataset, IP and Input, using BEDTools 2.17.0 (Quinlan and Hall, 2010) (genomcov -d -split -ibam) and removing all position with null coverage. Following the workflow described in (Schwartz et al., 2013) the Peak over the median (POM) was calculated by dividing median expression in the window by median expression of the gene, only considering exonic regions of genes. The sites with a POM score < 4 in the IP dataset were removed. The Peak Over Input (POI) score was then calculated by dividing POM score in the IP dataset by POM score in the Input dataset. Only the peak with POI > 2 were kept. MACS2 methylation sites detection was run (*-g 282000000 –nomodel* parameters) on bam files of IP datasets with the Input datasets serving for assessing the whole RNA distribution. Each of the 4 techniques was detecting small windows with potential methylation sites in very few datasets. To keep only robust methylation sites, the occurrence of each peak was assessed by counting in how many datasets a specific window is detected as a methylation site. Only windows found in 3 or more datasets were kept. BEDtools merge software (Quinlan and Hall, 2010) was run to regroup all overlapping windows with detected methylation sites. The overlapping region on their corresponding gene was searched: 5’UTR, 3’UTR, CDS, intron. The presence of the m^6^A consensus sequence RGACW in the sequence of the peak was assessed by calculating a motif score by adding score presence when one of this sequence was found: ‘GAACA’: 2, ‘GGACA’: 3, ‘GAACT’: 5, ‘GGACT’: 8. The union of all the methylation sites found by each of the four techniques was determined using BEDTools merge function. Two lists of potential methylation sites were then extracted: 86,418 methylation sites for cecum and 54,319 sites for liver tissue. With in-house python scripts we determined the overlap with the already detected m^6^A modification sites by MeRIP-Seq, CLIP-Seq, and Targets of m^6^A Readers, Erasers, and Writers using MeT-DB v2.0 database (Liu et al., 2018).

### Differential methylation analysis

Statistical analyzes were performed using the R package limma (Ritchie et al., 2015). Data were first normalized with TMM (Robinson and Oshlack, 2010) (edgeR package) and transformed with the voom (Law et al., 2014) function (limma package). Limma was then used to assess the statistical significance of observed differences in read counts. Three different linear models were derived to address three different questions. Differentially expressed genes between all pairs of the four conditions were first detected with *y ∼ BioCond + Sequencing + Library_batch*, where y is the normalized and transformed Input read counts (expression data), BioCond refers to the six biological conditions under study (CONV, GF, ex-GF, abx, vanco, Am). The sequencing batch was also accounted for through the *Sequencing* variable and the library preparation with *Library_batch* variable. Differentially expressed methylation sites were derived using a model on IP read counts to detect differential methylation (*y ∼ BioCond + Sequencing + Library_batch*). A third model (*y ∼ BioCond + Sequencing + Library_batch*) was applied on the calculated difference between IP and Input read counts. P-values resulting from the three models were adjusted for multiple comparisons using the BH procedure (Benjamini, 1995). Genes or methylation sites were considered statistically different when the adjusted p-value was lower than 0.05. The GUITAR plot was generated using the GUITAR package (Liu et al., 2018). KEGG pathway analysis was performed using the Enrichr tool (Chen et al., 2013; Kuleshov et al., 2016).

### qPCR from cecal content

gDNA was prepared from cecal content using the Power Soil DNA isolation kit (MoBio) following the manufacturer’s instructions. qPCR was performed as described (Reikvam et al., 2011) using EvaGreen Sso Fast Master Mix (Biorad), 500 nM primers and 50 ng input gDNA. Relative expression was determined using the ΔΔCT method with mpI genomic region as a reference as described (Reikvam et al., 2011).

### 16S DNA sequencing

16S DNA sequencing was performed as described previously. Briefly, the 16S rRNA gene amplification was performed by using the Nextflex 16s v1-v3 amplicon-seq kit. The 16S cecal content DNA was sequenced by using Illumina Miseq. Reads with a positive match with human, mice or phiX174 phage were removed. Library adapters, primer sequences, and base pairs occurring at 5’ and 3’ends with a Phred quality score <20 were trimmed off by using Alientrimmer (v0.4.0). Filtered high-quality reads were merged into amplicons with Flash (v1.2.11). Resulting amplicons were clustered into operational taxonomic units (OTU) with VSEARCH (v2.3.4)(Rognes et al., 2016) The process includes several steps for de-replication, singletons removal, and chimera detection. The clustering was performed at 97% sequence identity threshold, producing 1110 OTUs. The OTU taxonomic annotation was performed with the SILVA SSU (v128) database (Quast et al., 2013). The input amplicons were then mapped against the OTU set to get an OTU abundance table containing the number of reads associated with each OTU. All together, these stages and statistics are implemented in SHAMAN (shaman.pasteur.fr) (Quereda et al., 2016). The matrix of OTU count data was normalized at the OTU level by using the normalization method using total counts. Normalized counts were then summed within genera.

### QUANTIFICATION AND STATISTICAL ANALYSIS

All data are expressed as mean and standard error of the mean. The number of animals (n) for each group is indicated in the figure legends. Either student’s t-test or Ordinary one-way ANOVA for multiple comparisons were used for statistical analysis. This information is provided in the figure legends. For differential expression analysis p-values were adjusted using the Benjamini-Hochberg procedure as indicated in the figure legends. p-values < 0.05 were considered significant.

## SUPPLEMENTARY DATA

**Figure S1** m^6^A site analysis

**Figure S2** Gut microbiota composition

**Figure S3** Differential methylation compared to differential expression

**Figure S4** Differential methylation of Mat2a transcript in cecum

**Figure S5** LC-HRMS of m^6^A, m^1^A and m^5^C

**Fig. S1:**
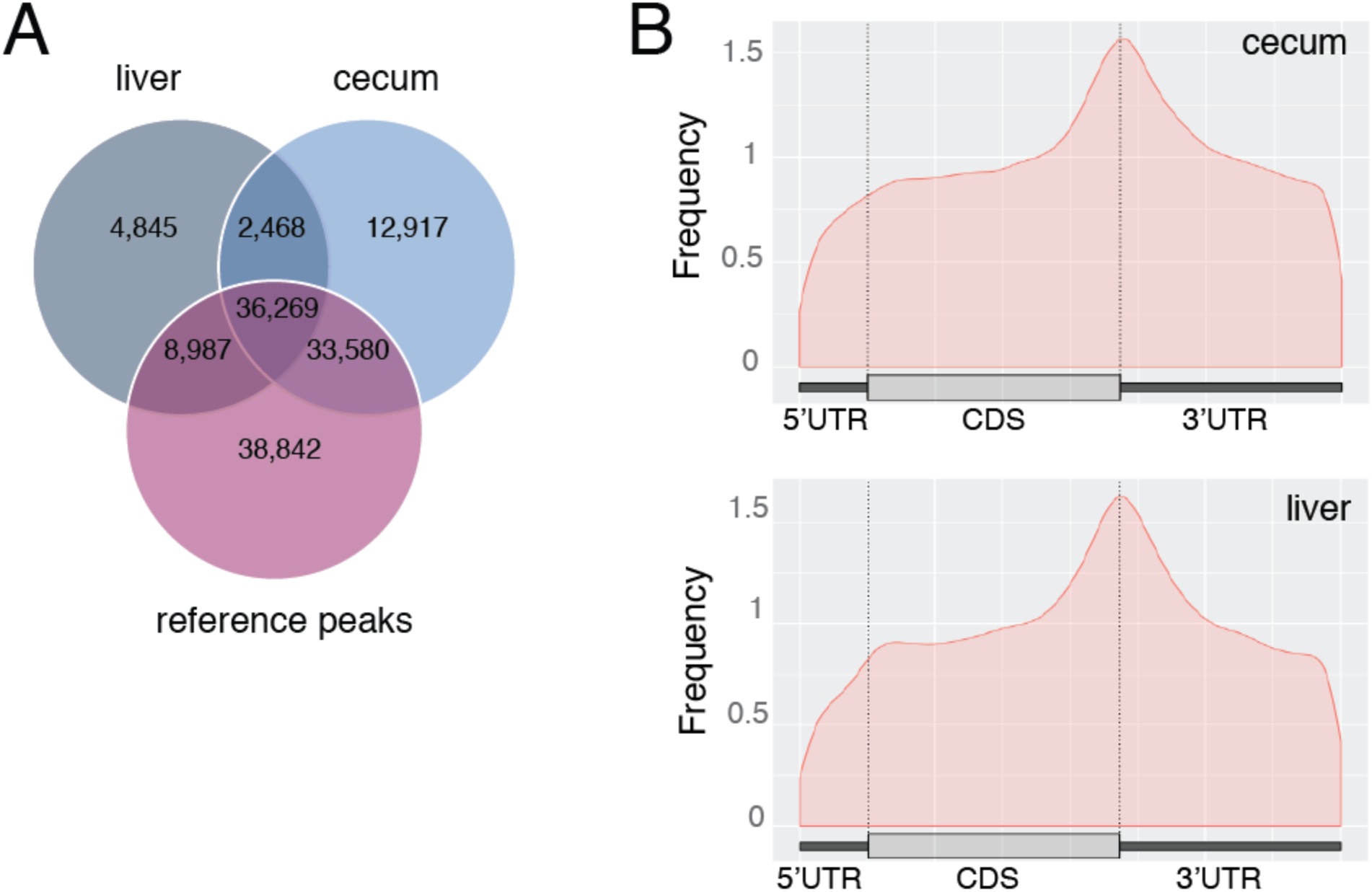
m^6^A site analysis, related to Figures 1 and 2. **A**, 46% (39,921/86,418) of m^6^A sites in the cecum are overlapping with at least one liver site, and 81% (69,849/86,418) of detected sites in the cecum are overlapping MeT-DB V2 reference site. For liver tissue, there are 74% (40,487/54,319) of methylation sites overlapping at least one site of cecal tissue, and 84% (45,807/54,319) overlapping MeT-DB V2 reference sites. The reference sites may be overlapping with several sites we detected (in average we found 2.36 and 2.2 m^6^A peaks per reference site for cecum and liver, respectively). **B**, Position of mapped m^6^A sites on the transcripts in liver and cecum determined using the GUITAR package.

**Fig. S2:**
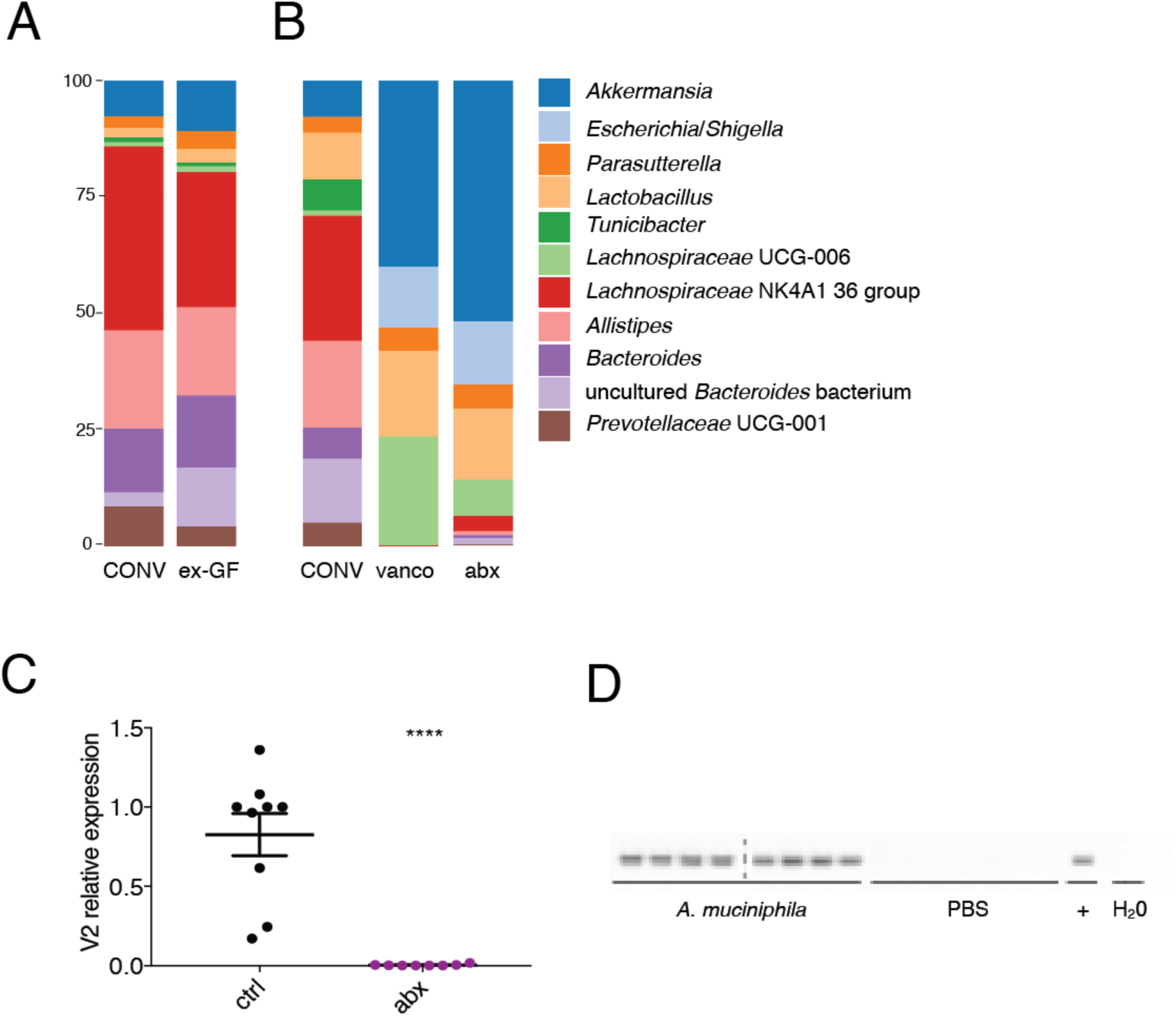
Gut microbiota composition, related to Figure 1 and 2. **A**, most abundant genera in cecal content of CONV (n=13) and ex-GF (n=16) mice colonized with CONV flora for 4 weeks by 16S DNA sequencing. Three independent experiments were performed; **B**, most abundant genera persisting in vancomycin/amphotericin B-treated mice (n= 12) and abx mice that had been treated with vancomycin, metronidazol, neomycin, ampicillin and amphotericin B every 12h for 21 days (n=10, 3 independent experiments); CONV n=12; **C,** quantitative PCR for V2 region of of cecal content from antibiotics-(abx, n=10) and vehicle treated (n=10) mice. Relative expression compared to mouse mpI genomic region. **** p-value < 0.001, unpaired t-test. **D**, *A*.*muciniphila-*specific PCR was performed on gDNA isolated from feces to control for colonization of GF mice.

**Fig. S3:**
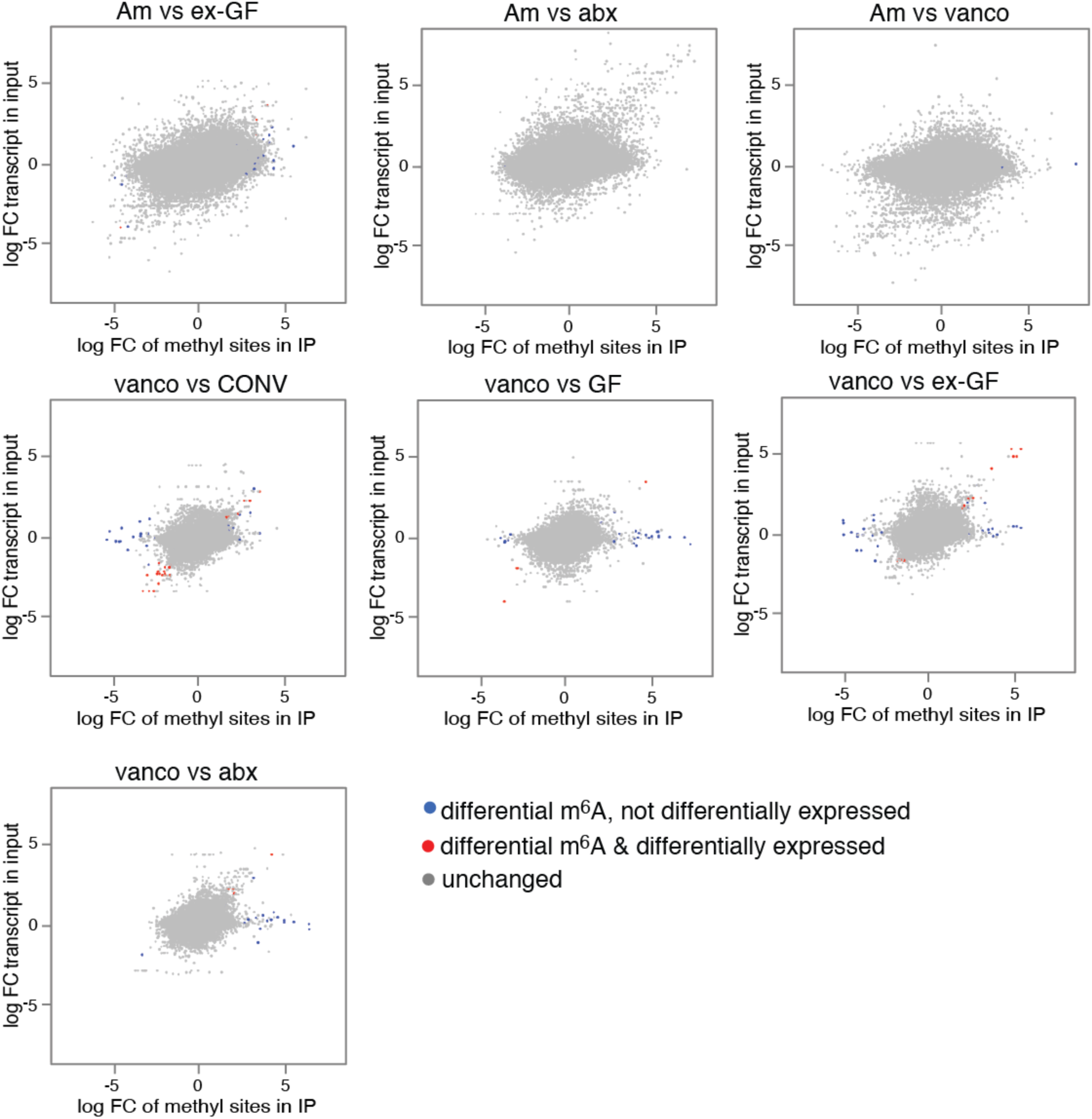
Differential methylation compared to differential expression, related to Figure 1. m^6^A-sites found to be differentially methylated compared to differential expression of transcripts in indicated conditions. Differentially methylated sites that are also differentially expressed on transcript level are in red, differentially methylated sites that are unchanged on transcript level, are in blue. Methylation sites that were not significantly changed are shown in grey. Cut-offs for differential expression are log fold change (FC) −1 to 1 and Benjamini-Hochberg-corrected p-values <0.05.

**Fig. S4:**
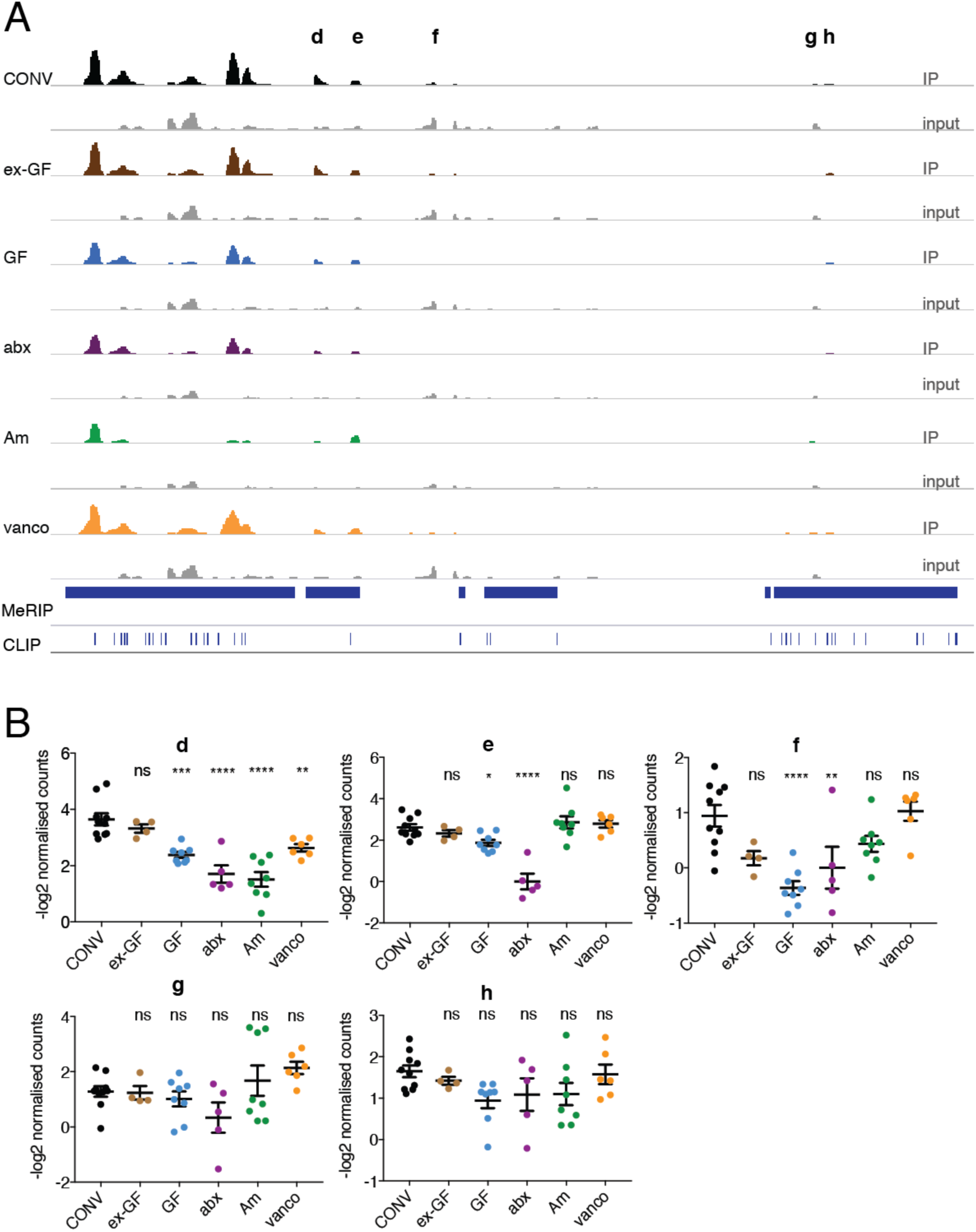
Differential methylation of Mat2a transcript in cecum, related to Figure 4. **A**, mean of read per million normalized reads of anti-m^6^A immunoprecipitates and input for Mat2a visualized using IGV. The MeDTB V2 database was used to indicate previously described methylation sites (MeRIP, blue bars) and sites of m^6^A-binding proteins identified using cross-linking immunoprecipitation (CLIP). **B**, quantification of select methylation sites (**d-h** from **A**). Shown is −log2 of normalized counts. Ordinary one-way ANOVA for multiple comparisons was performed. * p-value < 0.05; ** p-value < 0.01; *** p-value < 0.005; **** p-value < 0.001; ns, non-significant.

**Fig. S5:**
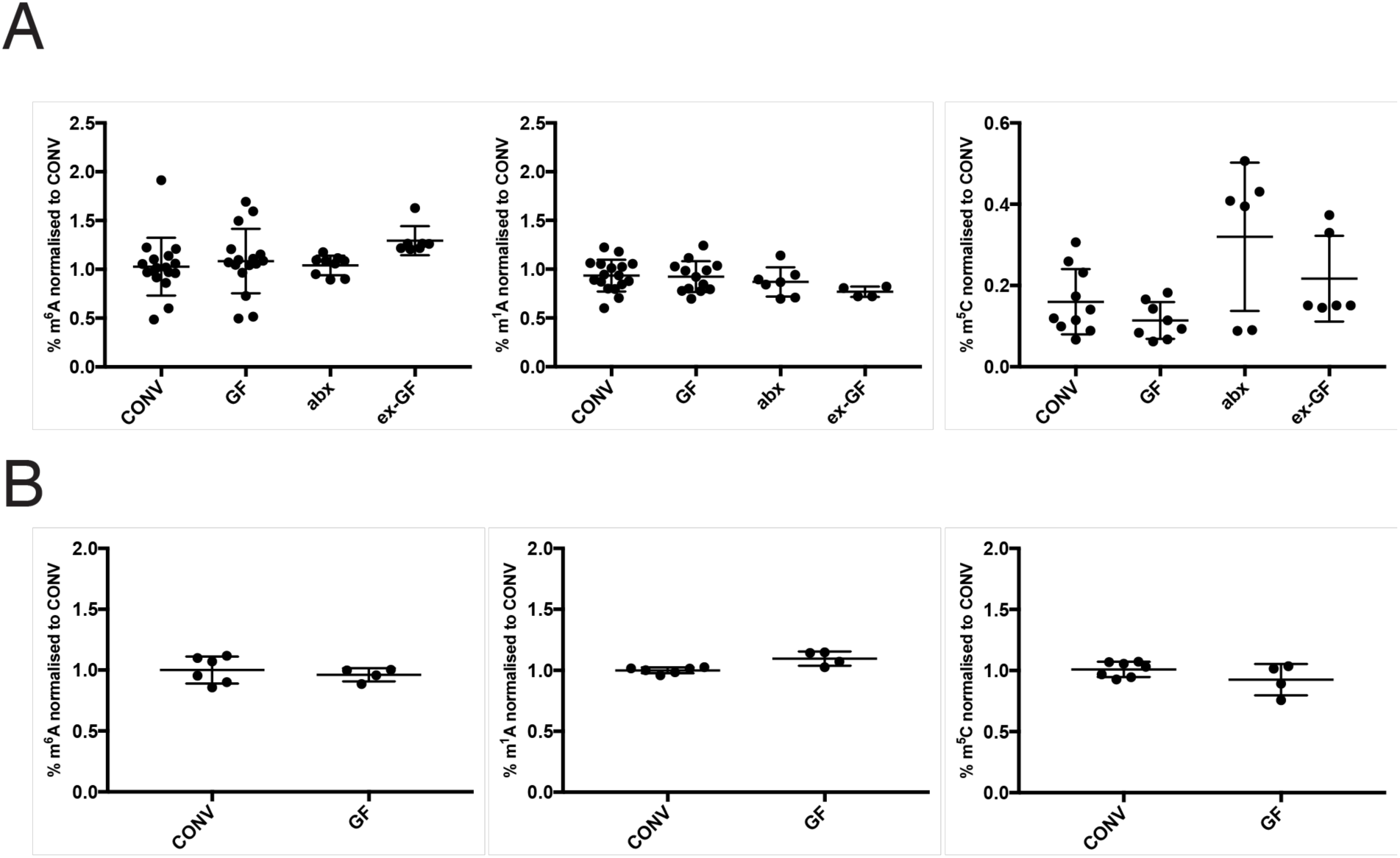
Relative quantification of selected mRNA modifications by liquid chromatography coupled to high resolution mass spectromtry (LC-HRMS) LC-HRMS of ribodepleted RNAs from **a**, cecum (m^6^A: CONV n=17, GF n=16, ex-GF n=7; abx n=9; m^1^A: CONV n=17, GF n=16, ex-GF n=4; abx n=7; m^5^C: CONV n=9, GF n=7, ex-GF n=6; abx n=7) and **b**, liver (CONV n=6; GF n=4); due to variability between MS analyses, samples for each batch (three in total) were normalised to the mean of the CONV values. Error bars depict SEM.

## References

Agus, A., Planchais, J., and Sokol, H. (2018). Gut Microbiota Regulation of Tryptophan Metabolism in Health and Disease. Cell Host & Microbe 23, 716–724.

Anders, S., Pyl, P.T., and Huber, W. (2015). HTSeq—a Python framework to work with highthroughput sequencing data. Bioinformatics 31, 166–169.

Bäckhed, F., Ding, H., Wang, T., Hooper, L.V., Koh, G.Y., Nagy, A., Semenkovich, C.F., and Gordon, J.I. (2004). The gut microbiota as an environmental factor that regulates fat storage. Proceedings of the National Academy of Sciences of the United States of America 101, 15718–15723.

Benjamini, Y.Y.H. (1995). Controlling the False Discovery Rate: A Practical and Powerful Approach to Multiple Testing. Journal of the Royal Statistical Society 57, 289–300.

Chen, E.Y., Tan, C.M., Kou, Y., Duan, Q., Wang, Z., Meirelles, G.V., Clark, N.R., and Ma’ayan, A. (2013). Enrichr: interactive and collaborative HTML5 gene list enrichment analysis tool. BMC Bioinformatics 14, 128.

Church, C., Lee, S., Bagg, E.A.L., McTaggart, J.S., Deacon, R., Gerken, T., Lee, A., Moir, L., Mecinović, J., Quwailid, M.M., et al. (2009). A Mouse Model for the Metabolic Effects of the Human Fat Mass and Obesity Associated FTO Gene. PLOS Genetics 5, e1000599.

Collado, M.C., Derrien, M., Isolauri, E., de Vos, W.M., and Salminen, S. (2007). Intestinal Integrity and Akkermansia muciniphila, a Mucin-Degrading Member of the Intestinal Microbiota Present in Infants, Adults, and the Elderly. Applied and Environmental Microbiology 73, 7767–7770.

Criscuolo, A., and Brisse, S. (2013). AlienTrimmer: A tool to quickly and accurately trim off multiple short contaminant sequences from high-throughput sequencing reads. Genomics 102, 500–506.

Derrien, M., Belzer, C., and de Vos, W.M. (2017). Akkermansia muciniphila and its role in regulating host functions. Microbial Pathogenesis 106, 171–181.

Dobin, A., Davis, C.A., Schlesinger, F., Drenkow, J., Zaleski, C., Jha, S., Batut, P., Chaisson, M., and Gingeras, T.R. (2013). STAR: ultrafast universal RNA-seq aligner. Bioinformatics 29, 15–21.

Dominissini, D., Moshitch-Moshkovitz, S., Salmon-Divon, M., Amariglio, N., and Rechavi, G. (2013). Transcriptome-wide mapping of N6-methyladenosine by m6A-seq based on immunocapturing and massively parallel sequencing. Nature Protocols 8, 176.

Dominissini, D., Moshitch-Moshkovitz, S., Schwartz, S., Salmon-Divon, M., Ungar, L., Osenberg, S., Cesarkas, K., Jacob-Hirsch, J., Amariglio, N., Kupiec, M., et al. (2012). Topology of the human and mouse m6A RNA methylomes revealed by m6A-seq. Nature 485, 201–206.

Ewels, P., Magnusson, M., Lundin, S., and Käller, M. (2016). MultiQC: summarize analysis results for multiple tools and samples in a single report. Bioinformatics 32, 3047–3048.

Fischer, J., Koch, L., Emmerling, C., Vierkotten, J., Peters, T., Brüning, J.C., and Rüther, U. (2009). Inactivation of the Fto gene protects from obesity. Nature 458, 894.

Frayling, T.M., Timpson, N.J., Weedon, M.N., Zeggini, E., Freathy, R.M., Lindgren, C.M., Perry, J.R.B., Elliott, K.S., Lango, H., Rayner, N.W., et al. (2007). A Common Variant in the FTO Gene Is Associated with Body Mass Index and Predisposes to Childhood and Adult Obesity. Science 316, 889–894.

Frye, M., and Blanco, S. (2016). Post-transcriptional modifications in development and stem cells. Development 143, 3871–3881.

Fustin, J.-M., Doi, M., Yamaguchi, Y., Hida, H., Nishimura, S., Yoshida, M., Isagawa, T., Morioka, Masaki S., Kakeya, H., Manabe, I., et al. (2013). RNA-Methylation-Dependent RNA Processing Controls the Speed of the Circadian Clock. Cell 155, 793–806.

Geula, S., Moshitch-Moshkovitz, S., Dominissini, D., Mansour, A.A., Kol, N., Salmon-Divon, M., Hershkovitz, V., Peer, E., Mor, N., Manor, Y.S., et al. (2015). m6A mRNA methylation facilitates resolution of naïve pluripotency toward differentiation. Science 347, 1002–1006.

Gokhale, Nandan S., McIntyre, Alexa B.R., McFadden, Michael J., Roder, Allison E., Kennedy, Edward M., Gandara, Jorge A., Hopcraft, Sharon E., Quicke, Kendra M., Vazquez, C., Willer, J., et al. (2016). N6-Methyladenosine in Flaviviridae Viral RNA Genomes Regulates Infection. Cell Host & Microbe 20, 654–665.

Heaver, S.L., Johnson, E.L., and Ley, R.E. (2018). Sphingolipids in host–microbial interactions. Current Opinion in Microbiology 43, 92–99.

Koh, A., De Vadder, F., Kovatcheva-Datchary, P., and Bäckhed, F. (2016). From Dietary Fiber to Host Physiology: Short-Chain Fatty Acids as Key Bacterial Metabolites. Cell 165, 1332– 1345.

Krautkramer, K.A., Kreznar, J.H., Romano, K.A., Vivas, E.I., Barrett-Wilt, G.A., Rabaglia, M.E., Keller, M.P., Attie, A.D., Rey, F.E., and Denu, J.M. (2016). Diet-Microbiota Interactions Mediate Global Epigenetic Programming in Multiple Host Tissues. Molecular Cell 64, 982– 992.

Kuleshov, M.V., Jones, M.R., Rouillard, A.D., Fernandez, N.F., Duan, Q., Wang, Z., Koplev, S., Jenkins, S.L., Jagodnik, K.M., Lachmann, A., et al. (2016). Enrichr: a comprehensive gene set enrichment analysis web server 2016 update. Nucleic Acids Research 44, W90–W97.

Law, C.W., Chen, Y., Shi, W., and Smyth, G.K. (2014). voom: precision weights unlock linear model analysis tools for RNA-seq read counts. Genome Biology 15, R29.

Li, H., Handsaker, B., Wysoker, A., Fennell, T., Ruan, J., Homer, N., Marth, G., Abecasis, G., and Durbin, R. (2009). The Sequence Alignment/Map format and SAMtools. Bioinformatics 25, 2078–2079.

Li, H.-B., Tong, J., Zhu, S., Batista, P.J., Duffy, E.E., Zhao, J., Bailis, W., Cao, G., Kroehling, L., Chen, Y., et al. (2017). m6A mRNA methylation controls T cell homeostasis by targeting the IL-7/STAT5/SOCS pathways. Nature 548, 338.

Lichinchi, G., Gao, S., Saletore, Y., Gonzalez, G.M., Bansal, V., Wang, Y., Mason, C.E., and Rana, T.M. (2016a). Dynamics of the human and viral m6A RNA methylomes during HIV-1 infection of T cells. Nature Microbiology 1, 16011.

Lichinchi, G., Zhao, B.S., Wu, Y., Lu, Z., Qin, Y., He, C., and Rana, T.M. (2016b). Dynamics of Human and Viral RNA Methylation during Zika Virus Infection. Cell Host & Microbe 20, 666–673.

Liu, H., Wang, H., Wei, Z., Zhang, S., Hua, G., Zhang, S.-W., Zhang, L., Gao, S.-J., Meng, J., Chen, X., et al. (2018). MeT-DB V2.0: elucidating context-specific functions of N6-methyladenosine methyltranscriptome. Nucleic Acids Research 46, D281–D287.

Liu, J., Yue, Y., Han, D., Wang, X., Fu, Y., Zhang, L., Jia, G., Yu, M., Lu, Z., Deng, X., et al. (2013). A METTL3–METTL14 complex mediates mammalian nuclear RNA N6-adenosine methylation. Nature Chemical Biology 10, 93.

Manes, N.P., Shulzhenko, N., Nuccio, A.G., Azeem, S., Morgun, A., and Nita-Lazar, A. (2017). Multi-omics Comparative Analysis Reveals Multiple Layers of Host Signaling Pathway Regulation by the Gut Microbiota. mSystems 2.

Mardinoglu, A., Shoaie, S., Bergentall, M., Ghaffari, P., Zhang, C., Larsson, E., Bäckhed, F., and Nielsen, J. (2015). The gut microbiota modulates host amino acid and glutathione metabolism in mice. Molecular Systems Biology 11.

Meyer, Kate D., Patil, Deepak P., Zhou, J., Zinoviev, A., Skabkin, Maxim A., Elemento, O., Pestova, Tatyana V., Qian, S.-B., and Jaffrey, Samie R. (2015). 5′ UTR m6A Promotes Cap-Independent Translation. Cell 163, 999–1010.

Meyer, Kate D., Saletore, Y., Zumbo, P., Elemento, O., Mason, Christopher E., and Jaffrey, Samie R. (2012). Comprehensive Analysis of mRNA Methylation Reveals Enrichment in 3′ UTRs and near Stop Codons. Cell 149, 1635–1646.

Mudge, J.M., and Harrow, J. (2015). Creating reference gene annotation for the mouse C57BL6/J genome assembly. Mammalian Genome 26, 366–378.

Nicholson, J.K., Holmes, E., Kinross, J., Burcelin, R., Gibson, G., Jia, W., and Pettersson, S. (2012). Host-Gut Microbiota Metabolic Interactions. Science 336, 1262–1267.

Peer, E., Rechavi, G., and Dominissini, D. (2017). Epitranscriptomics: regulation of mRNA metabolism through modifications. Current Opinion in Chemical Biology 41, 93–98.

Pendleton, K.E., Chen, B., Liu, K., Hunter, O.V., Xie, Y., Tu, B.P., and Conrad, N.K. (2017). The U6 snRNA m6A Methyltransferase METTL16 Regulates SAM Synthetase Intron Retention. Cell 169, 824–835.e814.

Pomerantz, S.C., and McCloskey, J.A. (1990). Analysis of RNA hydrolyzates by liquid chromatography-mass spectrometry. Methods in Enzymology 193, 796–824.

Quast, C., Pruesse, E., Yilmaz, P., Gerken, J., Schweer, T., Yarza, P., Peplies, J., and Glöckner, F.O. (2013). The SILVA ribosomal RNA gene database project: improved data processing and web-based tools. Nucleic Acids Research 41, D590–D596.

Quereda, J.J., Dussurget, O., Nahori, M.-A., Ghozlane, A., Volant, S., Dillies, M.-A., Regnault, B., Kennedy, S., Mondot, S., Villoing, B., et al. (2016). Bacteriocin from epidemic Listeria strains alters the host intestinal microbiota to favor infection. Proceedings of the National Academy of Sciences 113, 5706–5711.

Quinlan, A.R., and Hall, I.M. (2010). BEDTools: a flexible suite of utilities for comparing genomic features. Bioinformatics 26, 841–842.

Reikvam, D.H., Erofeev, A., Sandvik, A., Grcic, V., Jahnsen, F.L., Gaustad, P., McCoy, K.D., Macpherson, A.J., Meza-Zepeda, L.A., and Johansen, F.-E. (2011). Depletion of Murine Intestinal Microbiota: Effects on Gut Mucosa and Epithelial Gene Expression. PLOS ONE 6, e17996.

Ritchie, M.E., Phipson, B., Wu, D., Hu, Y., Law, C.W., Shi, W., and Smyth, G.K. (2015). limma powers differential expression analyses for RNA-sequencing and microarray studies. Nucleic Acids Research 43, e47–e47.

Robinson, M.D., and Oshlack, A. (2010). A scaling normalization method for differential expression analysis of RNA-seq data. Genome Biology 11, R25.

Rognes, T., Flouri, T., Nichols, B., Quince, C., and Mahé, F. (2016). VSEARCH: a versatile open source tool for metagenomics. PeerJ 4, e2584.

Schroeder, B.O., and Bäckhed, F. (2016). Signals from the gut microbiota to distant organs in physiology and disease. Nat Med 22, 1079–1089.

Schwartz, S., Agarwala, Sudeep D., Mumbach, Maxwell R., Jovanovic, M., Mertins, P., Shishkin, A., Tabach, Y., Mikkelsen, Tarjei S., Satija, R., Ruvkun, G., et al. (2013). High-Resolution Mapping Reveals a Conserved, Widespread, Dynamic mRNA Methylation Program in Yeast Meiosis. Cell 155, 1409–1421.

Schwartz, S., Mumbach, Maxwell R., Jovanovic, M., Wang, T., Maciag, K., Bushkin, G.G., Mertins, P., Ter-Ovanesyan, D., Habib, N., Cacchiarelli, D., et al. (2014). Perturbation of m6A Writers Reveals Two Distinct Classes of mRNA Methylation at Internal and 5’ Sites. Cell Reports 8, 284–296.

Shapiro, H., Kolodziejczyk, A.A., Halstuch, D., and Elinav, E. (2018). Bile acids in glucose metabolism in health and disease. The Journal of Experimental Medicine 215, 383–396.

Shima, H., Matsumoto, M., Ishigami, Y., Ebina, M., Muto, A., Sato, Y., Kumagai, S., Ochiai, K., Suzuki, T., and Igarashi, K. (2017). S-Adenosylmethionine Synthesis Is Regulated by Selective N6-Adenosine Methylation and mRNA Degradation Involving METTL16 and YTHDC1. Cell Reports 21, 3354–3363.

Sommer, F., and Bäckhed, F. (2016). Know your neighbor: Microbiota and host epithelial cells interact locally to control intestinal function and physiology. BioEssays 38, 455–464.

Suzuki, T., Ikeuchi, Y., Noma, A., Suzuki, T., and Sakaguchi, Y. (2007). Mass Spectrometric Identification and Characterization of RNA-Modifying Enzymes. In Methods in Enzymology (Academic Press), pp. 211–229.

Tan, B., and Gao, S.-J. (2018). RNA epitranscriptomics: Regulation of infection of RNA and DNA viruses by N6-methyladenosine (m6A). Reviews in Medical Virology 28, e1983.

Tirumuru, N., Zhao, B.S., Lu, W., Lu, Z., He, C., and Wu, L. (2016). N6-methyladenosine of HIV-1 RNA regulates viral infection and HIV-1 Gag protein expression. eLife 5, e15528.

Warda, A.S., Kretschmer, J., Hackert, P., Lenz, C., Urlaub, H., Höbartner, C., Sloan, K.E., and Bohnsack, M.T. (2017). Human METTL16 is a *N*^6^-methyladenosine (m^6^A) methyltransferase that targets pre-mRNAs and various non-coding RNAs. EMBO reports 18, 2004–2014.

Yoon, K.-J., Ringeling, F.R., Vissers, C., Jacob, F., Pokrass, M., Jimenez-Cyrus, D., Su, Y., Kim, N.-S., Zhu, Y., Zheng, L., et al. (2017). Temporal Control of Mammalian Cortical Neurogenesis by m6A Methylation. Cell 171, 877–889.e817.

